# Optimised FRET pairs and quantification approaches to detect the activation of Aurora kinase A at mitosis

**DOI:** 10.1101/562603

**Authors:** Giulia Bertolin, Florian Sizaire, Claire Déméautis, Catherine Chapuis, Fabienne Mérola, Marie Erard, Marc Tramier

## Abstract

Genetically-encoded Förster’s Resonance Energy Transfer (FRET) biosensors are indispensable tools to sense the spatiotemporal dynamics of signal transduction pathways. Investigating the crosstalk between different signalling pathways is becoming increasingly important to follow cell development and fate programs. To this end, FRET biosensors must be optimised to monitor multiple biochemical activities simultaneously and in single cells. In addition, their sensitivity must be increased to follow their activation even when the abundance of the biosensor is low.

We describe here the development of a second generation of Aurora kinase A/AURKA biosensors. First, we adapt the original AURKA biosensor –GFP-AURKA-mCherry– to multiplex FRET by using dark acceptors as ShadowG or ShadowY. Then, we use the novel superYFP acceptor protein to measure FRET by 2-colour Fluorescence Cross-Correlation Spectroscopy, in cytosolic regions where the abundance of AURKA is extremely low and undetectable with the original AURKA biosensor.

These results pave the way to the use of FRET biosensors to follow AURKA activation in conjunction with substrate-based activity biosensors. In addition, they open up the possibility of tracking the activation of small pools of AURKA and its interaction with novel substrates, which would otherwise remain undetectable with classical biochemical approaches.

## Introduction

Genetically-encoded Förster’s Resonance Energy Transfer (FRET) biosensors have revolutionised our knowledge of signal transduction pathways in the cell. The capability of sensing the activation of kinases, the activity of caspases, or the transport of second messengers as Ca^2+^ or cAMP opened up the possibility of following biochemical reactions in real time and with a spatiotemporal resolution (Sizaire and Tramier, 2017; Greenwald et al., 2018; Palmer et al., 2011). The novel frontier of these probes consists in combining two or more FRET biosensors at a time to unravel the interdependence of signal transduction pathways, an approach known as multiplex FRET (Piljic and Schultz, 2008; Carlson and Campbell, 2009; Ai et al., 2008; Ding et al., 2011; Su et al., 2013; Demeautis et al., 2017; Ross et al., 2018).

AURKA is a serine/threonine kinase with multiple functions in the cell. AURKA was first described to be a mitotic protein with multiple partners and playing key roles in centrosome maturation, in the regulation of mitotic timing, and in building and stabilising the mitotic spindle (Nikonova et al., 2013). Nonetheless, it is now becoming increasingly clear that AURKA has several roles outside of mitosis, such as favouring neurite outgrowth (Mori et al., 2009), transcriptional activity through the *MYC* promoter (Zheng et al., 2016), and mitochondrial homeostasis (Bertolin et al., 2018; Grant et al., 2018). In this light, it is still unknown whether mitotic and non-mitotic functions of AURKA are made possible by a single, recycling pool of the kinase or by different pools at specific subcellular locations. Given the fact that the abundance of AURKA is regulated throughout the cell cycle, sensitive techniques are required to decipher the role of AURKA at selected compartments and especially when the abundance of the protein is low as in G1 phase.

We previously engineered an AURKA FRET biosensor where the donor-acceptor FRET pair flanks the entire AURKA sequence. This new strategy for FRET biosensors allowed us to replace the endogenous kinase in mammalian cells, and to report on its activation at centrosomes both at mitosis and during G1 phase (Bertolin et al., 2016). We here develop two independent strategies to ameliorate this probe: first, a single-colour AURKA biosensor for multiplex FRET and second, a method to estimate FRET efficiency in regions where the abundance of AURKA is low. We show that FRET donor-acceptor pairs with dark acceptors are preferable for engineering multiplex FRET approaches with the AURKA biosensor, rather than fluorescence anisotropy. We also optimise 2-colour Fluorescence Correlation Spectroscopy (2c-FCCS), together with an improved donor-acceptor FRET pair, to detect AURKA activation in cytoplasmic regions with low AURKA abundance.

## Results and discussion

### Homo-FRET within the AURKA biosensor is undetectable by fluorescence polarisation microscopy

We previously generated an AURKA FRET biosensor (GFP-AURKA-mCherry) reporting on the autophosphorylation of the protein kinase AURKA on Thr288 (Bertolin et al., 2016). Biochemical studies reported that this post-translational modification primes the kinase for activation towards a given substrate (Bayliss et al., 2003; Cheetham, 2002; Zhang et al., 2007). In the perspective of using our FRET biosensor in a multiplex FRET context, we modified the GFP-AURKA-mCherry biosensor to create a single-colour FRET sensor to detect homo-FRET. We replaced the mCherry acceptor protein located at the carboxy terminus of the original AURKA biosensor with an Enhanced Green Fluorescent Protein (mEGFP), thereby creating a GFP-AURKA-GFP biosensor (Fig. 1A). Changes in the conformation of GFP-AURKA-GFP were analysed by following the fluorescence anisotropy of the biosensor, obtained by fluorescence polarization microscopy (Tramier and Coppey-Moisan, 2008). With this approach, decreased anisotropy would correspond to FRET within the biosensor.

**Fig. 1.**
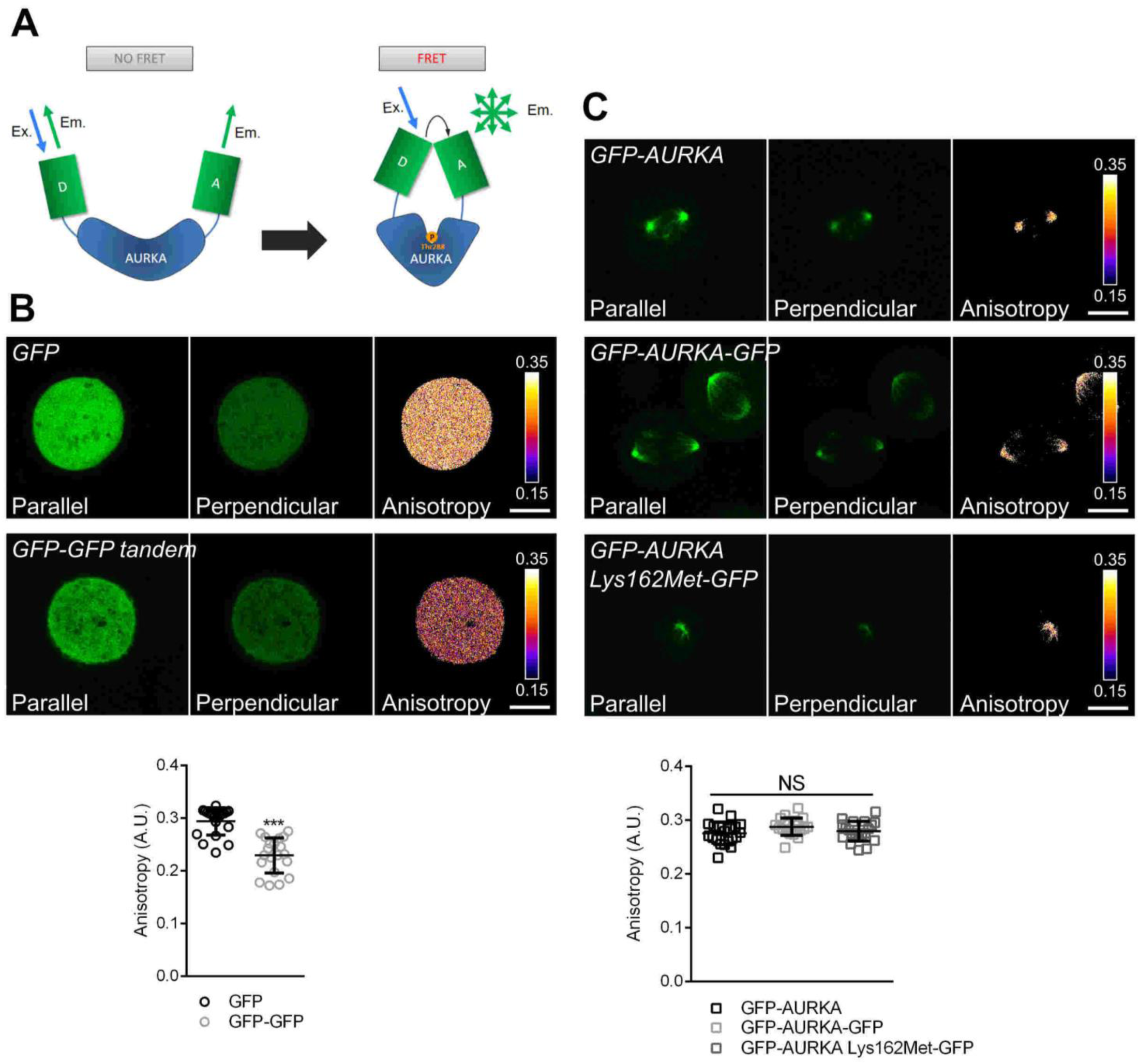
The GFP-AURKA-GFP biosensor does not allow the detection of homo-FRET. **A.** Model illustrating the mode of action of the GFP-AURKA-GFP biosensor. This biosensor switches from an open-to-close conformation upon autophosphorylation of AURKA on Thr288. While no depolarisation is observed in the open conformation, the closed conformation of the biosensor induces a partial depolarisation of the acceptor, due to the FRET phenomenon. Of note, the real three-dimensional orientation of the two GFPs is unknown. **B.** Representative parallel and perpendicular polarised images, and corresponding anisotropy quantification of U2OS cells synchronised at mitosis and expressing a GFP monomer (upper panels) or a GFP tandem dimer (lower panels). **C.** Representative parallel and perpendicular polarised images, and corresponding anisotropy quantification of U2OS cells synchronised at mitosis and expressing GFP-AURKA, GFP-AURKA-GFP or GFP-AURKA Lys162Met-GFP localised at the mitotic spindle. The pseudocolor scale in **B.** and **C.** represents pixel-by-pixel anisotropy. Transfection conditions are indicated in italics. Anisotropy values for individual cells are represented as circles in **B.**, and as squares in **C.** Data represent means ± s.d. of one representative experiment of three. *n*=30 cells per condition. Scale bar: 10 nm. \*\*\**P*<0.001 against the ‘GFP’ condition; NS: not significant.

We synchronised U2OS cells at mitosis to benefit from a cell cycle phase in which AURKA is active and highly abundant at the mitotic spindle (Nikonova et al., 2013), and where the GFP-AURKA-mCherry biosensor displays significant hetero-FRET (Bertolin et al., 2016). To establish the maximum decrease in anisotropy detectable with our fluorophore pair, we compared a GFP monomer and a tandem dimer. For both constructs, we imaged the parallel and perpendicular components of the emitted fluorescence. This was performed by exciting the sample with a polarised laser, followed by the detection through an analyser in a confocal microscope setup. Similarly to what previously described, the dimer displays a decreased anisotropy (mean anisotropy: 0.22) compared to the GFP monomer (mean anisotropy: 0.30) in mitotic cells, thus indicating the homotransfer between the two GFP fluorophores harbouring different orientations (Fig. 1B) (Tramier and Coppey-Moisan, 2008). This is consistent with the fact that the rotation period of the GFP fluorophore is slower than fluorescence lifetime (Gautier et al., 2001; Swaminathan et al., 1997). We then evaluated the anisotropy of GFP-AURKA, of the biosensor GFP-AURKA-GFP and of the kinase-dead version of the biosensor, GFP-AURKA Lys162Met-GFP (Bertolin et al., 2016), on the mitotic spindle. As previously described (Bertolin et al., 2016), the Lys162Met variant of AURKA induces mitotic spindle abnormalities such as monopolar or multipolar spindles. The anisotropy of GFP-AURKA was not different from the one of the GFP monomer, indicating that the presence of the kinase does not perturb the anisotropy of GFP *per se* (Fig. 1C). However, we did not detect any significant difference between the anisotropy of GFP-AURKA and of the biosensor, both in the activated and in the kinase-dead version (Fig. 1C).

Although fluorescence anisotropy was recently shown to be a suitable approach for FRET biosensors (Ross et al., 2018), relative differences in anisotropy between active and inactive biosensors are often very low. In the case of the AURKA biosensor, the difference between the active and the inactive forms of the biosensor was not significant, with the GFP-AURKA-GFP showing a slightly increased anisotropy rather than a decreased one. This could be due to the clustering of the GFP-AURKA-GFP biosensor at the spindle. Indeed, the high mechanical tension taking place at this compartment at mitosis could keep the fluorophores in a stable conformation, altering the rotational motility of GFP. Alternatively, GFP rotation could also be perturbed by the presence of selected interactors of AURKA at the spindle, which could maintain the biosensor in a fixed conformation. Last, our results could also indicate that the two GFP fluorophores are in a parallel conformation, making differences in anisotropy undetectable and preventing the use of this approach for multiplex FRET in our biosensor setup.

### The dark acceptors ShadowG and ShadowY allow for single-colour FRET/FLIM measurements

In the perspective of optimising a multiplex FRET strategy for the AURKA biosensor, we then explored whether dark acceptors as ShadowG (Murakoshi et al., 2015) or ShadowY (Murakoshi and Shibata, 2017) could be used as efficient alternatives to fluorescence anisotropy to this end. These proteins were previously demonstrated to be non-fluorescent variants of the Yellow Fluorescent Protein (YFP) preventing spectral bleedthrough of the first acceptor in the second donor channel, thereby resulting in a suitable strategy for multiplex FRET design. As shown in the original reports, they also behave as good acceptors in FRET/FLIM pairs for mGFP or mClover, respectively. These properties of ShadowG and ShadowY allow for multi-colour imaging and FRET/FLIM studies with other fluorescent proteins, and we previously took advantage of these properties of ShadowG for the setup of multiplex FRET analyses. In particular, ShadowG turned out to be an efficient acceptor for the cyan fluorescent protein mTFP1 as well (Demeautis et al., 2017), and we used the mTFP1-ShadowG donor-acceptor pair in conjunction with a second one constituted by the Large Stoke Shift fluorescent protein mOrange (LSSmOrange) and mKate2. This combination of fluorophores used in FRET biosensors allowed us to record two biochemical activities simultaneously (Demeautis et al., 2017; Ringer et al., 2017).

In light of these previous data, we first monitored the effect of using ShadowG and ShadowY as dark acceptors for the cyan donor protein mTurquoise2. We observed a decrease in the lifetime of the donor of ~500 psec when ShadowG was present in a tandem construct, and of ~800 psec when ShadowY was used (Supplementary Fig. 1A), demonstrating that both fluorophores are excellent acceptors for mTurquoise2. We then replaced the GFP-mCherry donor-acceptor pair of the original AURKA biosensor with mTurquoise2 and ShadowG or ShadowY, thereby creating ShadowG-AURKA-mTurquoise2 or ShadowY-AURKA-mTurquoise2 (Fig. 2B). We then tested the conformational changes of these two biosensors by FRET/FLIM in U2OS cells synchronised at mitosis, by calculating the net difference in the lifetime (ΔLifetime) between the donor-only construct (AURKA-mTurquoise2) and ShadowG-AURKA-mTurquoise2 or ShadowY-AURKA-mTurquoise2 at the mitotic spindle. ΔLifetime was used to standardise the decrease in lifetime due to FRET, as lifetime variation is the measured parameter directly dependent on the standard deviation of the experimental setup used (50 psec with our setup (Bertolin et al., 2016)). We measured a mean ΔLifetime of ~150 psec for both biosensors (Fig. 2B-C), which was similar to the decrease observed for the original GFP-AURKA-mCherry one (Bertolin et al., 2016). With the same approach, we also tested the impact of kinase-dead AURKA on FRET efficiencies. ΔLifetime values for both biosensors carrying the Lys162Met mutation were half than their wild-type counterparts, indicating lower FRET efficiencies when AURKA is catalytically inactive. However, it must be noted that the difference in lifetime between normal and kinase-dead AURKA biosensors with dark acceptors (~50 psec) was less remarkable than what observed with the GFP/mCherry AURKA biosensor (~120 psec), where the GFP-AURKA Lys162Met-mCherry lifetime was comparable to the one of the donor alone (Bertolin et al., 2016).

**Fig. 2.**
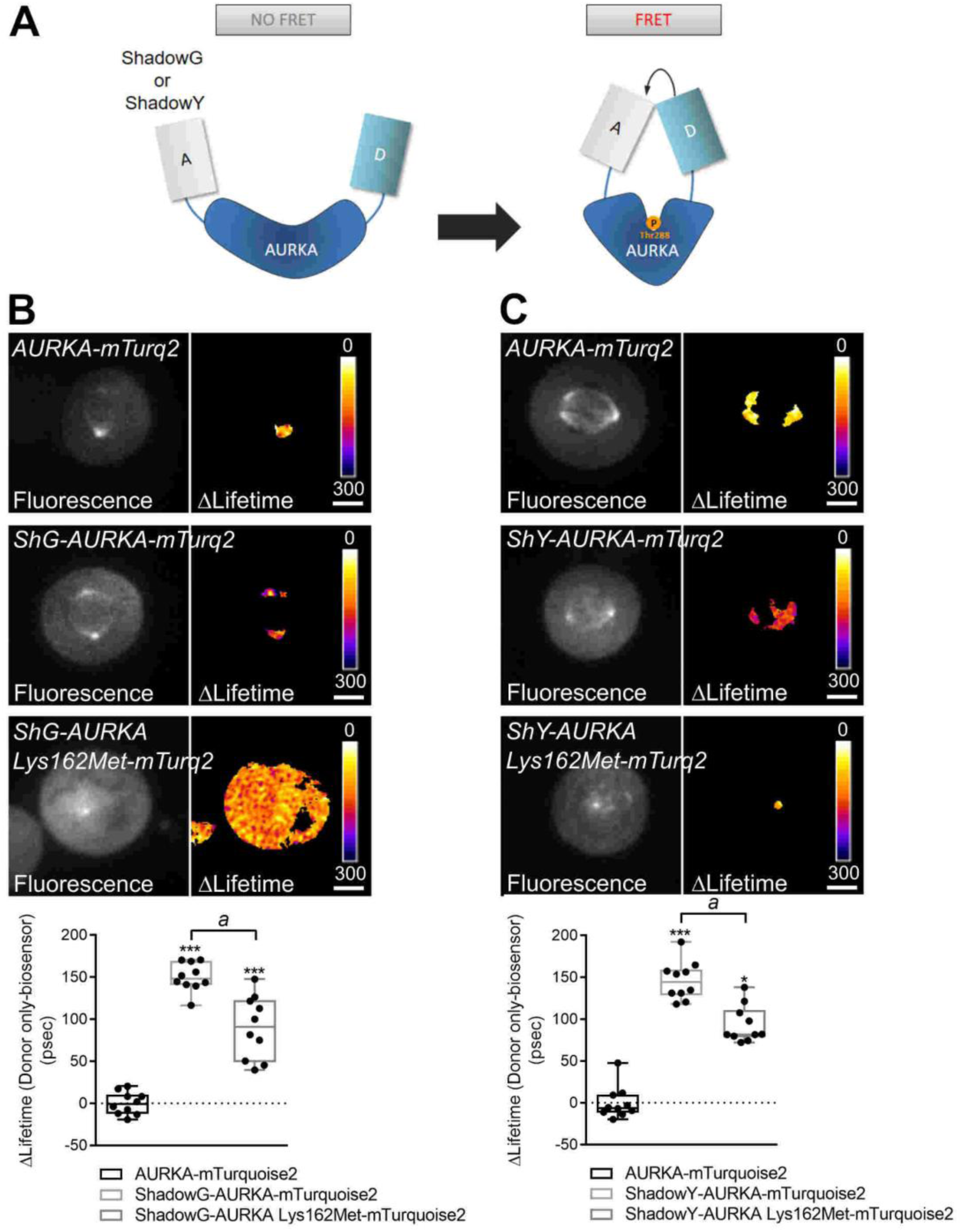
ShadowG and ShadowY are efficient dark acceptors for mTurquoise2 in the AURKA biosensor. **A.** Model illustrating the mode of action of the ShadowG-AURKA-mTurquoise2 or the ShadowY-AURKA-mTurquoise2 biosensors. These biosensors switch from an open-to-close conformation upon autophosphorylation of AURKA on Thr288, bringing the donor and the acceptor in vicinity and allowing FRET detection**. B. and C.** (Upper panels) Representative fluorescence (mTurquoise2 channel) and ΔLifetime (donor only-biosensor) images of U2OS cells expressing the indicated constructs and synchronised at mitosis. (Lower panel) Corresponding ΔLifetime quantification at the mitotic spindle. ShG: ShadowG; ShY: ShadowY; mTurq2: mTurquoise2. ΔLifetime values for individual cells are represented as black dots in each boxplot. The bar in boxplots represents the median; whiskers extend from the 10th to the 90th percentiles. *n*=10 cells per condition of one representative experiment (of three). Scale bar: 10 nm. \*\*\**P*<0.001 against the ‘AURKA-mTurquoise2’ condition; ^*a*^*P*<0.001 compared to the ‘ShadowG-AURKA-mTurquoise2’ condition in **B.** or ^*a*^*P*<0.05 compared to the ‘ShadowY-AURKA-mTurquoise2’ condition in **C.**

Our results show that the replacement of the donor-acceptor pair from GFP-mCherry to mTurquoise2-ShadowG or ShadowY maintains FRET efficiency. In addition, both dark biosensors show a lowered FRET efficiency when the kinase is catalytically dead, although this difference is less pronounced than in the original GFP-AURKA-mCherry biosensor. It should also be noted that the performance of a FRET biosensor cannot be predicted from tandem constructs. In spite of the significantly better performance of the ShadowY-mTurquoise2 tandem fluorescence anisotropy for the design of multiplex FRET experiments with the AURKA kinase biosensor. In this light, cumulating a biosensor of AURKA containing a dark acceptor with a substrate-based biosensor of the kinase flanked by the LSSmOrange/mKate2 donor-acceptor pair could allow the simultaneous monitoring of AURKA activation by auto-phosophorylation on Thr288 and its activity towards a substrate, on the same cell and with a spatiotemporal resolution.

### The GFP-AURKA-mCherry biosensor is not suitable to estimate FRET by two-colour FCCS

It is known that AURKA is a multifunctional kinase with several subcellular locations, with both mitotic and non-mitotic functions. While mitotic AURKA is highly abundant in cells (Nikonova et al., 2013), the low abundance of non-mitotic pools of AURKA often represent a limitation in exploring their function. Our current approach based on FRET/FLIM allows to analyse the activation of the kinase only where it is particularly abundant (i.e. the centrosome, the mitotic spindle or mitochondria) (Bertolin et compared to the ShadowG-mTurquoise2 one (Supplementary Fig. 1A), the corresponding AURKA biosensors show an identical response (Fig. 2B and C). Therefore, two additional parameters should be kept into consideration. First, the nature of the protein probed within the biosensor and second, the possibility that the conformational changes of this protein fall within the Förster’s radius of the donor-acceptor pair or not.

However, the use of biosensors containing dark acceptors should be considered as a suitable alternative to al., 2016, 2018), possibly excluding from the analysis other cytosolic pools of AURKA too scarce to be analysed with this microscopy technique. Therefore, we turned to two-colour Fluorescence Cross-Correlation Spectroscopy (2c-FCCS) to estimate FRET in regions of the cytoplasm where the abundance of AURKA is low and the number of photons is insufficient to estimate the activation of AURKA by fluorescence lifetime.

In 2c-FCCS, the fluctuating signal intensities of the red and the green channels in and out of a confocal volume are monitored in real time, and their auto-correlation function is computed together with a cross-correlation function between the two channels. Besides these measurements, we used the temporal correlation of each fluorophore excited by different lasers (pulsed or continuous) to remove the spectral bleed-through of GFP into the mCherry channel. This was done following a previously published strategy (Padilla-Parra et al., 2014, 2011) based on Fluorescence Lifetime Correlation Spectroscopy (FLCS) (Böhmer et al., 2002). We excited GFP with a pulsed laser and mCherry with a continuous laser. We then gave a specific weight to each photon in the green and in the red channel using pulsed or continuous calculated lifetime filters. In order to eliminate cross-talk effects between GFP and mCherry, these weights were used to build the auto-correlation curves for each fluorophore, and the cross-correlation curve. Applying this strategy to two separate molecules of GFP and mCherry gives a null cross-correlation, thereby eliminating potential artefacts in 2c-FCCS experiments due to spectral bleed-through (Padilla-Parra et al., 2014, 2011). In 2c-FCCS, the presence of FRET is indicated by a simultaneous decrease of the amplitude of the cross-correlation curve and the increased amplitude of the green auto-correlation curve (Foo et al., 2012; Kohl et al., 2002). This allows to use 2c-FCCS to estimate FRET in co-diffusing molecules, as it is the case for FRET biosensors.

With this approach, we estimated the green and red auto-correlation curves of the original AURKA biosensor (GFP-AURKA-mCherry) and of the kinase-dead GFP-AURKA Lys 162Met-mCherry variant in cells synchronised at mitosis. For each cell analysed, we measured three randomly-chosen points in the cytosol to estimate FRET efficiency in this compartment. We did not detect any significant difference between active and kinase-dead AURKA when we compared the ratio of the amplitudes of the red or green auto-correlation curves with the amplitude of the cross-correlation curve (Fig. 3). This was in apparent contradiction with previous data acquired in cells where the cytosol was strong enough to perform FRET/FLIM analyses (Bertolin et al., 2016), and where we detected FRET with the GFP-AURKA-mCherry biosensor. Therefore, we concluded that the ratio between the cross-correlation and the green auto-correlation is not sufficiently robust to detect changes due to FRET in the original AURKA biosensor.

**Fig. 3.**
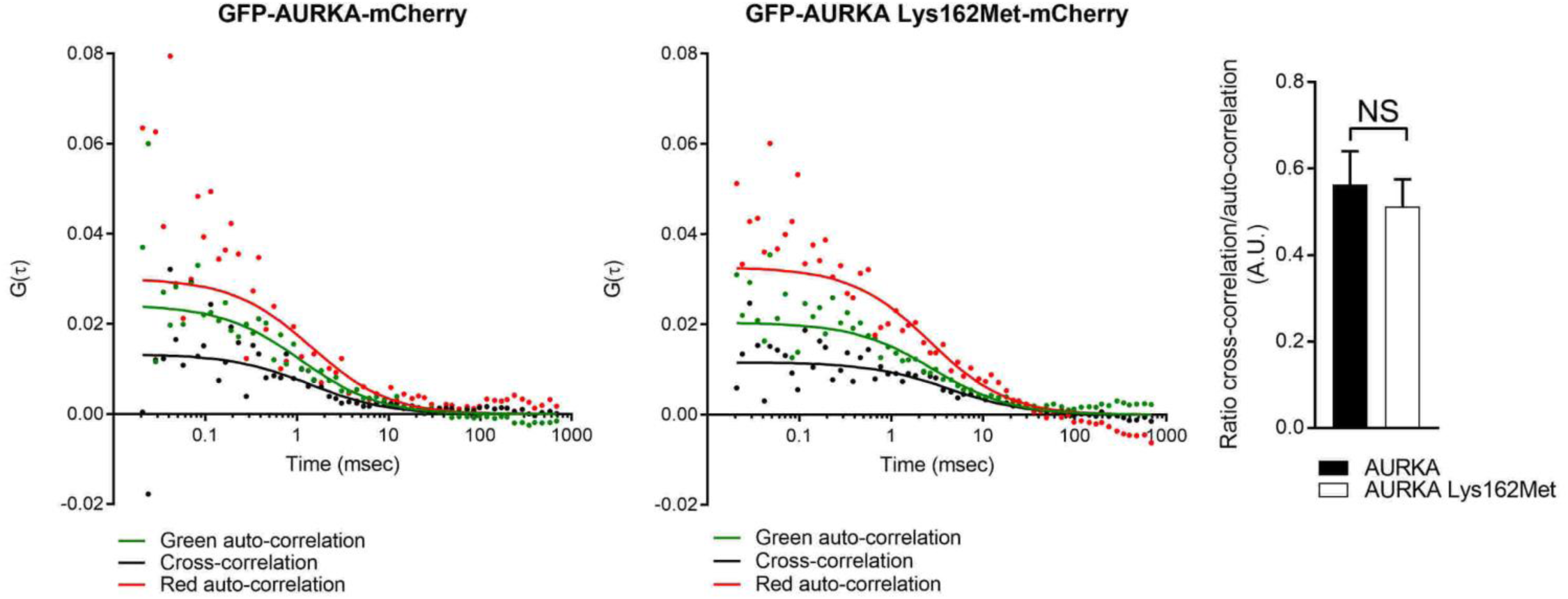
The GFP-AURKA-mCherry biosensor does allow FRET detection by 2c-FCCS. Green and red auto-correlation curves, together with the respective cross-correlation curves issued from one representative U2OS cell expressing GFP-AURKA-mCherry (left panel) or GFP-AURKA Lys162Met-mCherry (middle panel) and synchronised at mitosis. Measurements were taken in the cytosol; one independent point per condition is shown. Experimental values obtained for each curve are represented as dots. Time is expressed in msec. (Right panel) Ratio of the cross-correlation/green auto-correlation values for GFP-AURKA-mCherry (black bar) and GFP-AURKA Lys162Met-mCherry (white bar). Each *n* represents the average of three independent points per cell at 0.01 msec; n=10 cells per condition of one representative experiment (of three). NS: not significant.

As FRET cannot be estimated by 2c-FCCS using the original AURKA biosensor, it is possible that mCherry represents a limiting factor for the measurement of FRET via 2c-FCCS, due to its longer dark state compared to the one of EGFP-like proteins (Dean et al., 2011), and its lower brightness (15.84 for mCherry *vs* 33.6 for mEGFP; www.fpbase.org). Changes in the donor-acceptor fluorescent pair should then be envisaged to adapt the AURKA biosensor to 2c-FCCS, choosing an acceptor with a higher brightness to ameliorate the robustness of the amplitude of the different correlation curves.

### Replacing the donor-acceptor pair is mandatory to adapt the AURKA biosensor to two-colour FCCS

To find a suitable donor-acceptor pair to adapt the AURKA biosensor to 2c-FCCS, we first searched for a bright acceptor. We decided to replace the usual acceptors of cyan fluorescent proteins Citrine/Venus (brightness of approximately 69; www.fpbase.org), by superYFP, a novel dimeric yellow fluorescent protein with high brightness and based on dLanYFP (the dLanYFP brightness is 112.5; www.fpbase.org); the full characterisation of the physicochemical properties of superYFP will be described elsewhere. We coupled superYFP to mTurquoise2, as this appeared to be an efficient donor fluorophore in the dark versions of the AURKA biosensor (Fig. 2). We then tested the capacity of mTurquoise2 to act as a donor of energy for superYFP in a tandem construct. mTurquoise2 displayed a ΔLifetime of ~600 psec when superYFP was present in U2OS cells synchronised at mitosis (Supplementary Fig. 1B), indicating that this donor-acceptor couple is suitable for FRET/FLIM analyses.

In light of these data, we then engineered the superYFP-AURKA-mTurquoise2 biosensor (Fig. 4A). To validate this new biosensor, we evaluated its conformational changes at the mitotic spindle by FRET/FLIM. Again, we compared the ΔLifetime between the donor-only construct AURKA-mTurquoise2 and superYFP-AURKA-mTurquoise2, and we observed a mean ΔLifetime of ~210 psec in the presence of the acceptor (Fig.4B). Unlike the GFP-AURKA-mCherry original biosensor, the presence of the AURKA Lys162Met mutation did not abolish FRET efficiency, as observed for the variants of the AURKA biosensor containing dark acceptors. The differences in Förster’s radius (R0) for the mTurquoise/superYFP pair (59 Å) compared to GFP/mCherry (51 Å; (Albertazzi et al., 2009)) could explain the behaviour of the two biosensors. Applying the FRET equation with the usual value κ^2^=2/3, the estimated distance between donor and acceptor is of 83 Å for the normal, active AURKA, and more than 97 Å for the kinase-dead. Using the same distances for the superYFP-AURKA-mTurquoise2 active and dead biosensors, we calculated lifetime values (2.97 ns for the active form; 3.18 ns for the kinase-dead) compatible with what we measured experimentally (3.15 ns for the active; 3.25 ns for the kinase-dead), taking 3.35 ns as lifetime of the donor alone. It should be noted that the time-gated detection of our system does not yield an unequivocal absolute lifetime value for mTurquoise2 (Theoretical lifetime of mTurquoise2: 4 ns (Goedhart et al., 2012); lifetime measured with our system: 3.35 ns). Nevertheless, we recently provided evidence that net differences between donor-only and biosensors constructs are maintained (Demeautis et al., 2017). Taken together, the difference between active and kinase-dead form is the main parameter to be taken into account to estimate the quality of any AURKA biosensor. In this light, the difference in the mTurquoise/superYFP biosensors is of 100 psec, whereas the one observed using the GFP/mCherry biosensors is of 130 psec (Bertolin et al., 2016). This indicates that when FRET is calculated by FLIM, this new biosensor does not show a better performance.

**Fig. 4.**
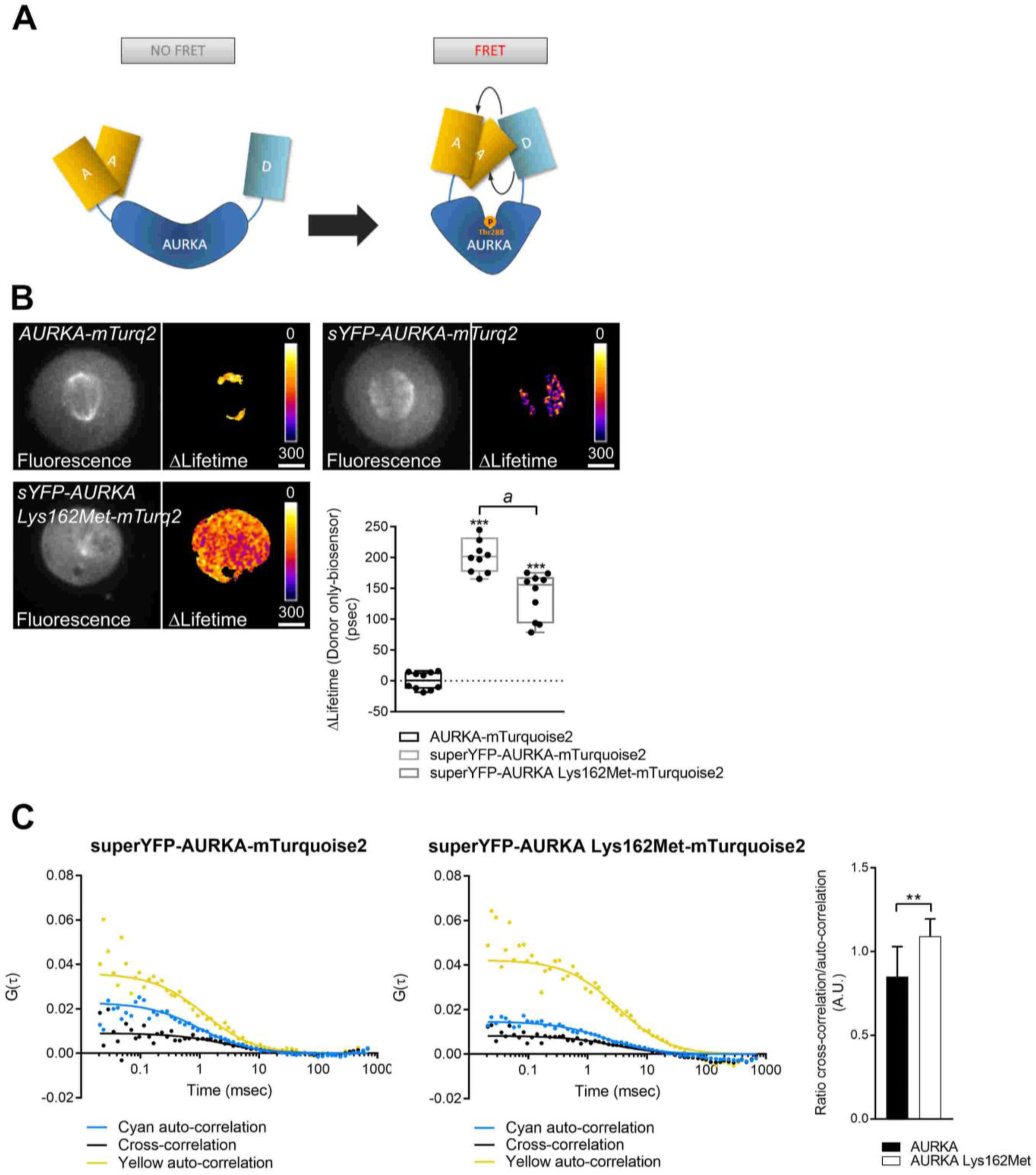
superYFP coupled to mTurquoise2 ameliorates FRET/FLIM detection and allows FRET detection by 2c-FCCS. **A**. Model illustrating the mode of action of the superYFP-AURKA-mTurquoise2 biosensor. It switches from an open-to-close conformation upon autophosphorylation of AURKA on Thr288, and allowing FRET measurements. Of note, superYFP is a dimeric acceptor. **B.** Representative fluorescence (mTurquoise2 channel) and ΔLifetime (donor only-biosensor) images of U2OS cells expressing the indicated constructs and synchronised at mitosis, together with the corresponding ΔLifetime quantification at the mitotic spindle. sYFP: superYFP; mTurq2: mTurquoise2. ΔLifetime values for individual cells are represented as black dots in each boxplot. The bar in boxplots represents the median; whiskers extend from the 10th to the 90th percentiles. *n*=10 cells per condition of one representative experiment (of three). Scale bar: 10 µm. **C.** Cyan and yellow auto-correlation curves, together with the respective cross-correlation curves issued from one representative U2OS cell expressing superYFP-AURKA-mTurquoise2 (left panel) or superYFP-AURKA Lys162Met-mTurquoise2 (middle panel) and synchronised at mitosis. Measurements were taken in the cytosol; one independent point per condition is shown. Time is expressed in msec. Experimental values obtained for each curve are represented as dots. (Right panel) Ratio of the cross-correlation/cyan auto-correlation values for superYFP-AURKA-mTurquoise2 (black bar) and superYFP-AURKA Lys162Met-mTurquoise2 (white bar). Each *n* represents the average of three independent points per cell at 0.01 msec; *n* =10 cells per condition of one representative experiment (of three) \*\*\**P* <0.001 against the ‘AURKA-mTurquoise2’ condition (**B.**), \*\**P* <0.01 against the ‘AURKA’ condition (**C.**); ^*a*^*P*<0.001 compared to the ‘superYFP-AURKA-mTurquoise2’ condition (**B.**)

However, we used this version of the AURKA biosensor for 2c-FCCS, potentially benefiting from the greater brightness of superYFP in the computing of the cross-correlation curve. We calculated the cyan and yellow auto-correlation curves of superYFP-AURKA-mTurquoise2 and of superYFP-AURKA Lys162Met-mTurquoise2 in U2OS cells synchronised at mitosis (Fig. 4C). As done previously for the GFP-AURKA-mCherry biosensor, we measured 3 independent points per cell, chosen in a random manner in the cytosol. We observed a simultaneous increase of the cyan auto-correlation curve and a decrease of the cross-correlation curve in cells expressing the superYFP-AURKA-mTurquoise2 biosensor when compared to the kinase-dead one. The ratio between the cross-correlation and the cyan auto-correlation was lower in cells expressing the active biosensor than in cells expressing the kinase-dead one (Fig. 4C, right panel).

We also noticed that the amplitude of the auto-correlation curve of superYFP is higher than the one of mTurquoise2. We reasoned that this could be linked to some photophysical properties of superYFP, induced by the laser power used for monitoring its diffusion. We thus monitored the amplitude of the auto-correlation curve of superYFP after modifying the power of the 514 nm laser. Given that the 2c-FCCS measurements were done with a laser power of ~ 1% (Fig. 4C), we chose four different laser powers for comparison: 0.5%, 1%, 3% and 5%. We didn’t observe any significant alteration in the yellow amplitude, which undergoes a 20% increase at most when switching the 514 nm laser from 0.5% to 1% and then remains constant between 1%, 3% and 5% (Supplementary Fig. 2). The fact that there is no significant change in the amplitude of the yellow auto-correlation curve when changing the laser power suggests that a modification of the photophysical behaviour of superYFP may not be the responsible of the observed higher amplitude. Another possibility is that the high amplitude of the yellow auto-correlation curve in 2c-FCCS analyses reflects a lower concentration of the yellow moiety in comparison to the cyan one. Given that the superYFP derives from dLanYFP, this could be due to the slower maturation of superYFP compared to mTurquoise2.

Taken together, the variations in amplitude of the cyan auto-correlation and of the cross-correlation curves meet the requirements for FRET within the superYFP-AURKA-mTurquoise2 biosensor. Although correlation analyses were proven to be a useful approach to characterise single-molecule FRET (Nguyen et al., 2012; Nettels et al., 2015), we here report for the first time the use of 2c-FCCS to monitor FRET within a genetically-encoded biosensor. Last, this new version of the AURKA biosensor containing an mTurquoise/superYFP donor-acceptor FRET pair is a suitable tool to investigate the activation of discrete pools of AURKA with low abundance and with high spatiotemporal resolution.

## Conclusions

Here we propose several improvements to the original AURKA FRET biosensor, both in terms of changing the donor-acceptor FRET pair for multiplex analyses, or in exploring new approaches to estimate FRET efficiency. It might be useful to point out that although fluorescence anisotropy was successfully used to follow multiplex FRET among PKA, ERK and cAMP (Ross et al., 2018), this is clearly not a suitable approach for the AURKA biosensor. The results here described indicate that there is no “golden rule” for FRET biosensors, and that the capacity of every biosensor to give a satisfactory response with a particular methodology approaches must be carefully evaluated.

Changing the donor-acceptor FRET pairs was mandatory to cumulate the analysis of the conformational changes of AURKA and of its enzymatic activity towards a particular substrate. Given the fact that AURKA is a multifunctional kinase involved in several biochemical pathways (Nikonova et al., 2013), designing a strategy for multiplex FRET is a first, mandatory step to start understanding how AURKA auto-activates and interacts with its wide range of substrates. The spatiotemporal resolution AURKA activation and its concomitant interaction with a substrate, or more at a time, is a crucial aspect in the future development of therapeutic strategies in epithelial cancers showing an overexpression of AURKA.

Changes in the way FRET efficiency is calculated, together with improvements in the donor-acceptor FRET pair, were also required to evaluate the activation of AURKA when its abundance is low. 2c-FCCS with the mTurquoise/superYFP pair efficiently responds to this need, allowing to monitor the activation of AURKA in the cytosol. We previously described that AURKA is activated at mitochondria (Bertolin et al., 2018), a compartment where the kinase is abundant enough to perform FRET/FLIM analyses. The mTurquoise/superYFP AURKA FRET biosensor used in conjunction with fluorescent markers for other subcellular compartments could potentially shed light in the activation of AURKA at other locations, where the kinase is less abundant. Such activation could lead to the identification of novel roles of AURKA, together with novel substrates.

## Materials and Methods

### Expression vectors and molecular cloning procedures

The GFP-AURKA, GFP-AURKA-mCherry and the GFP-AURKA Lys162Met-mCherry vectors suitable for expression in mammalian cells were previously described (Bertolin et al., 2016). To obtain GFP-AURKA-GFP, GFP-AURKA under the *AURKA* minimal promoter sequence (CTTCCGG) was subcloned into a pCDNA6 vector (Thermo Fisher Scientific) into the Eam1105I/NheI restriction sites, and a second EGFP was inserted into the NheI/NotI sites. The GFP monomer and tandem dimer were previously described (Tramier and Coppey-Moisan, 2008). A plasmid containing the mTurquoise2 was obtained from Dorus Gadella, together with the plasmids encoding ShadowG and ShadowY, which were obtained from Hideji Murakoshi. These plasmids were obtained via Addgene and have been previously described (Murakoshi et al., 2015; Murakoshi and Shibata, 2017; Mastop et al., 2017). To obtain AURKA-mTurquoise2, AURKA was subcloned into a pEGFP-N1 vector into the NotI/BamHI cloning site, while EGFP was replaced by mTurquoise2. To obtain ShadowG-AURKA-mTurquoise2, ShadowY-AURKA-mTurquoise2 and superYFP-AURKA-mTurquoise2, ShadowG, ShadowY or superYFP were inserted into the XhoI/HindIII restriction sites of AURKA-mTurquoise2, respectively. To obtain the mTurquoise2-ShadowG tandem, a tandem containing mTFP1 and ShadowG (Demeautis et al., 2017) was digested with BglII/AgeI to replace mTFP1 with mTurquoise2. To obtain the mTurquoise2-ShadowY tandem, a pEGFP-N1 backbone vector containing mTurquoise2 was digested with BglII/BamHI to insert ShadowY. To create the mTurquoise2-superYFP tandem, a pEGFP-N1 backbone vector containing a tandem of Aquamarine and superYFP was digested with AgeI/BglII to replace Aquamarine with mTurquoise2. To obtain a vector mTurquoise2 alone, the mTurquoise2-superYFP tandem was digested with BamHI to eliminate the superYFP fluorophore. Cloning reactions were performed with the NEBuilder HiFi DNA Assembly Mix (New England Biolabs) or by T4 DNA ligase (Thermo Fisher Scientific). All cloning reactions were verified on a 3130 XL sequencer (Applied Biosystems). All restriction enzymes were purchased from Thermo Fisher Scientific. The Lys162Met variant for all AURKA biosensors analysed in the study were obtained from the corresponding wild-type constructs by QuikChange site-directed mutagenesis (Stratagene) with the following primers: 5’-CAAGTTTATTCTGGCTCTTATGGTGTTATTTAAAGCTCA GCT-3’ (sense) and 5’-AGCTGAGCTTTAAATAACACCATAAGAGCCAGAATAA ACTTG-3’ (anti-sense).

### Cell culture and synchronisation procedures

U2OS cells free from mycoplasma were purchased from American Type Culture Collection (ATCC, HTB-96) and were grown in Dulbecco’s modified Eagle’s medium (DMEM, Sigma-Aldrich) supplemented with 10% fetal bovine serum (Life Technologies, Thermo Fisher Scientific), 1% L-glutamine (Life Technologies, Thermo Fisher Scientific) and 1% penicillin– streptomycin (Life Technologies, Thermo Fisher Scientific). The generation of GFP-AURKA, GFP-AURKA-mCherry and GFP-AURKA Lys 162Met-mCherry stable cell lines was previously described (Bertolin et al., 2016). GFP-AURKA-GFP and GFP-AURKA Lys162Met-GFP stable clones cells were generated by transfecting U2OS cells with X-tremeGENE HP transfection reagent (Roche), following the manufacturer’s indications. Stable clones were selected in DMEM supplemented with 10% fetal bovine serum, 1% L-glutamine, 1% penicillin–streptomycin and 500 µg/ml Geneticin (Invivogen). Stable and transient transfections were performed with X-tremeGENE HP transfection reagent (Roche) or with Lipofectamine 2000 (Life Technologies, Thermo Fisher Scientific), according to the manufacturer’s instructions. For live microscopy, cells were incubated in phenol red-free Leibovitz’s L-15 medium (Thermo Fisher Scientific), supplemented with 20% fetal bovine serum, 1% L-glutamine and 1% penicillin–streptomycin. Mitotic cells were obtained after synchronisation at the G2/M transition with 100 ng/ml nocodazole (Sigma-Aldrich) for 16 h. Cells were washed twice with 1X PBS and incubated with prewarmed imaging medium for 30 min to reach metaphase before performing fluorescence anisotropy, FRET/FLIM or 2c-FCCS analyses. All live microscopy experiments were performed at 37°C in Nunc Lab-Tek II Chamber slides (Thermo Fisher Scientific).

### Fluorescence polarisation microscopy

Fluorescence polarisation analyses were performed on a SP8 (Leica) inverted confocal microscope with a 63X oil immersion objective (NA 1.4), and using the Leica Acquisition Suite (LAS)-AF software. As the excitation light coming out of the laser is naturally polarized, an analyser integrated in the Leica emission filter wheel was placed in the emission pathway before the pinhole. The analyser rotates from a parallel to a perpendicular orientation to perform a sequential acquisition of the two corresponding polarised images. Images of GFP polarisation were acquired using a 488 nm argon laser and emission was selected between 500 and 550 nm. Image analysis was performed using the Fiji (ImageJ) software and fluorescence anisotropy was calculated as in (Tramier and Coppey-Moisan, 2008), using the following equation: *r* = (*I*_∥_ -*I*_⊥_)/(*I*_∥_ + 2*I*_⊥_), where *r* is the fluorescence anisotropy, *I*_∥_and *I*_⊥_ are the parallel and the perpendicular polarisations and *I*_∥_ + 2*I*_⊥_ is the total polarisation-independent emission.

### FLIM microscopy

FLIM analyses were performed in the time domain with a time-gated custom-built setup as described in (Bertolin et al., 2016) and driven by the Inscoper hardware (Inscoper). Briefly, cells were excited at 440+/-10 nm and at 480+/-10 nm using a white light laser and emission was selected using a band pass filter of 483/35 nm and of 525/50 nm for mTurquoise2 and GFP, respectively. Fluorescence lifetime was calculated with five sequential temporal gates of 2.2 nsec each. Mean pixel-by-pixel lifetime was calculated using the following equation: ⟨τ⟩=ΣΔti⋅Ii/ΣIi where Δti is the delay time of the *i*th image acquired following a laser pulse, and I is the pixel-by-pixel fluorescence intensity in each image. Lifetime measurements and online calculations were performed with the Inscoper software (Inscoper). Pixel-by-pixel lifetime was calculated only when fluorescence intensity was above 3000 grey levels. To calculate ΔLifetime, the mean lifetime of the cells expressing the donor alone (AURKA-mTurquoise2) was calculated for each independent experiment. This mean lifetime value was then used to normalise data in all the analysed conditions.

### Two-colour fluorescence cross-correlation spectroscopy

2c-FCCS analyses were carried out on a Leica a SP8 (Leica) inverted confocal microscope with a 63X water objective with correcting ring (NA 1.2), combined with a PicoQuant time-correlated single photon counting (TCSPC) module and a Fluorescence Lifetime Correlation Spectroscopy (FLCS) integration (PicoHarp, PicoQuant). For the GFP/mCherry donor-acceptor pair the following laser sources were used: a pulsed one at 470 nm and a continuous one at 561 nm, while for the mTurquoise2/superYFP pair we used a pulsed laser source at 440 nm and a continuous one at 514 nm. Fluorescence emission was collected simultaneously using two avalanche photodiode detectors (APD and Tau-SPAD, PicoQuant). The two emission channels used were, 500-550 nm and 581-654 nm for GFP/mCherry and 467-499 nm and 525-565 nm for mTurquoise2/superYFP. 2c-FCCS measurements were carried out with the Symphotime software (PicoQuant), integratd into the LAS-AF software (Leica). For each cell, the fluctuation of the fluorescence intensity for both donor-acceptor pairs (green/red or cyan/yellow) in the confocal volume was analysed by monitoring three independent points in the cytosol. These points were analysed sequentially and in non-photobleaching conditions for 30 sec each. Auto-correlation and cross-correlation curves were reconstructed from the fluorescence decay with the Symphotime software and the lifetime integration as previously reported (Padilla-Parra et al., 2014, 2011), with the Symphotime software. Data were fit using the 3D diffusion model, with one diffusing species (Padilla-Parra et al., 2014, 2011). The ratio between cross-correlation and green (for GFP/mCherry) or cyan (for mTurquoise2/superYFP) auto-correlation was used to estimate FRET efficiency.

### Statistical analyses

One-way ANOVA and the Dunnet’s method were used to compare the effects of the transfected AURKA plasmids on GFP fluorescence anisotropy (Fig. 1C). One-way ANOVA and the Tukey’s method were used to compare the effect of the transfected vectors on fluorescence lifetime (Fig. 2B; Supplementary Fig. 1A). One-way ANOVA on ranks and the Kruskal-Wallis method were used to compare the effect of the transfected vectors on fluorescence lifetime (Fig. 2C, 4B).The Student’s t-test was used to compare the effect of a GFP monomer and a tandem dimer on fluorescence anisotropy (Fig. 1B) or to compare the effects of superYFP on the lifetime of mTurquoise2 (Supplementary Fig. 1B); the Mann-Whitney test was used to compare the cross-correlation/auto-correlation ratio between normal and Lys162Met AURKA (Fig. 3 and 4C).

**Supplementary Fig. 1.**
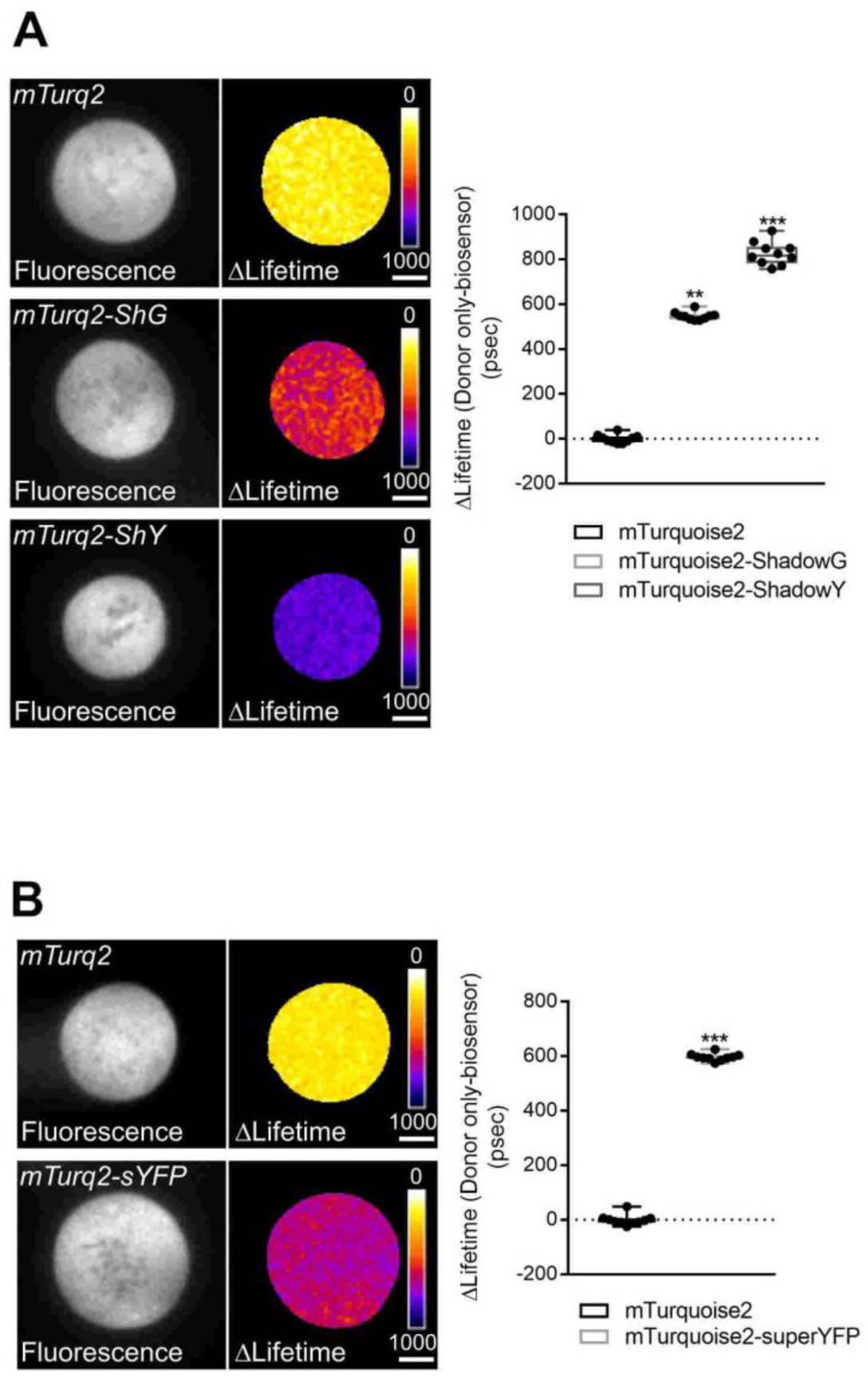
mTurquoise2-ShadowG, ShadowY or superYFP are efficient donor-acceptor FRET pairs. **A. and B.** (Left panels) Representative fluorescence (mTurquoise2 channel) and ΔLifetime (donor only-biosensor) images of U2OS cells expressing the indicated constructs and synchronised at mitosis. ShG: ShadowG; ShY: ShadowY; sYFP: superYFP; mTurq2: mTurquoise2. (Right panel). ΔLifetime values for individual cells represented as black dots in each boxplot. The bar in boxplots represents the median; whiskers extend from the 10th to the 90th percentiles. *n* =10 cells per condition of one representative experiment (of three). Scale bar: 10 nm. \*\**P* <0.01 and \*\*\**P* <0.001 against the ‘mTurquoise2’ condition.

**Supplementary Fig. 2.**
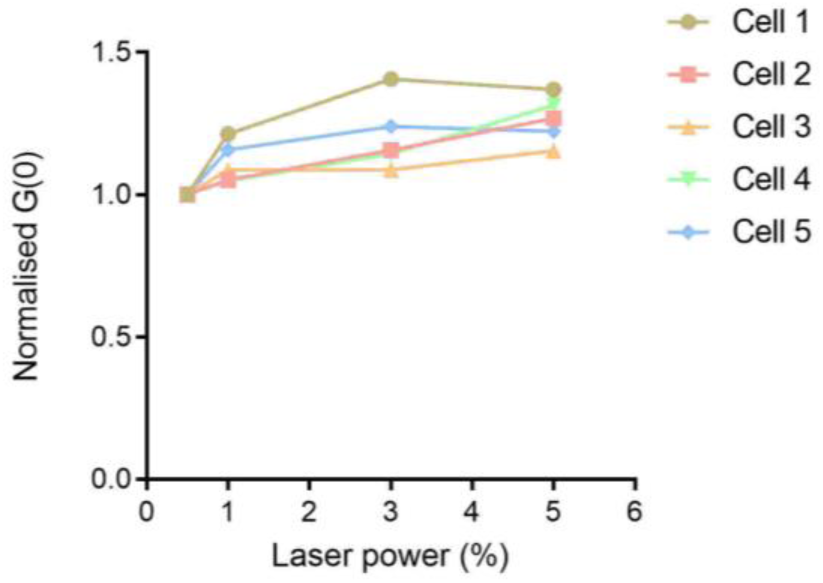
The amplitude of superYFP is not altered by increasing the 514 nm laser power. Normalised G(0) values obtained from five independent U2OS cells expressing the superYFP-AURKA-mTurquoise2 FRET biosensor and synchronised at mitosis. Cells were illuminated with the 514 nm laser only at increasing powers, i.e. 0.5%, 1%, 3% and 5%. Measurements were taken in the cytosol; each point represents the average of three independent points per cell at 0.01 msec.

## Author contributions

G.B. designed, performed and analysed the experiments, wrote the manuscript and provided funding; F.S. and C.D. performed and analysed the experiments, CC provided technical help; M.E. and F.M. provided the superYFP coding sequence and relative vectors, shared unpublished data and provided advice; M.T. conceptualised the study, coordinated the work and provided funding.

## Acknowledgements

We thank all the members of the Microscopy-Rennes Imaging Center (Biologie, Santé, Innovation Technologique, BIOSIT, Rennes, France) for assistance and in particular X. Pinson for providing an ImageJ macro for the analysis of fluorescence anisotropy. We also thank Laurent Deleurme from the Flow cytometry and Cell Sorting platform (BIOSIT, Rennes, France) for assistance, together with A. Webb and S. Ku for technical help. This work was supported by the Comité Nationale de la Recherche Scientifique, by the Ligue Contre le Cancer Comité d’Ille et Vilaine et Comité des Côtes d’Armor to G.B. and by the Comité d’Ille et Vilaine, Comité du Maine et Loire et Comité de la Sarthe to MT, and by the *Infrastructures en Biologie Santé et Agronomie* (IBiSA), région Bretagne and Rennes Métropole for the development of the technology for rapid FLIM measurements. FS was supported by a fellowship from Région Bretagne and the University of Rennes1, together with an additional fellowship from the *Ligue Nationale Contre le Cancer*.

